# Glutamate-Weighted Magnetic Resonance Imaging (GluCEST) Detects Effects of Transcranial Magnetic Stimulation to the Motor Cortex

**DOI:** 10.1101/2021.11.18.469133

**Authors:** Abigail T.J. Cember, Benjamin L. Deck, Apoorva Kelkar, Olu Faseyitan, Jared P. Zimmerman, Brian Erickson, Mark A. Elliott, H. Branch Coslett, Roy H. Hamilton, Ravinder Reddy, John D. Medaglia

## Abstract

**Transcranial magnetic stimulation (TMS) is used in several FDA-approved treatments** and, increasingly, to treat neurological disorders in off-label uses. However, the mechanism by which TMS causes physiological change is unclear, as are the origins of response variability in the general population. Ideally, objective *in vivo* biomarkers could shed light on these unknowns and eventually inform personalized interventions. **Continuous theta burst stimulation (cTBS)** is a form of TMS which has been observed to reduce motor evoked potentials (MEPs) for 60 minutes or longer post-stimulation, although the consistency of this effect and its mechanism continue to be under debate. Here, we use **glutamate-weighted chemical exchange saturation transfer (gluCEST) magnetic resonance imaging (MRI)** at ultra-high magnetic field (7T) to measure changes in glutamate concentration at the site of cTBS. We find that gluCEST signal in the ipsilateral hemisphere of the brain generally decreases in response to cTBS, whereas consistent changes were not detected in the contralateral or in subjects receiving a sham stimulation.

**One Sentence Summary:** We used glutamate-weighted Chemical Exchange Saturation Transfer (GluCEST) imaging to detect changes in glutamate contrast in the brains of young, healthy adults undergoing transcranial magnetic stimulation (TMS) to the motor cortex.

## Introduction

Transcranial magnetic stimulation (TMS) is a non-invasive brain stimulation technique which has been employed extensively by cognitive neuroscientists, physiologists, and clinicians^1^. TMS uses electromagnetic induction to generate electrical current in the cerebral cortex, and causes depolarization of neurons at the site of stimulation. Specifically, TMS uses a strong and rapidly fluctuating electrical current which is transmitted through loops of conductive wires surrounded by a protective casing and held in close proximity to the skull. The magnetic field generated by the current penetrates the scalp and skull, inducing electrical and subsequent chemical changes within the cortex^2,3^. TMS to the motor cortex (M1) has been shown to activate cortical interneurons and pyramidal neurons, leading to corticospinal tract activation and, ultimately, muscle contraction^4^. When applied to other regions of the cortex, TMS can interrupt normal cognitive function and can improve the condition of those with complex neuropsychiatric disorders such as depression^5,6^. Given the already wide-ranging application of this technique, it is ever more imperative for scientists and clinicians to understand the mechanisms by which TMS influences the brain and subsequent behaviors.

To this end, multiple TMS protocols have been developed. The focus of our study is a particular approach known as continuous theta burst stimulation (cTBS)^7^, consisting of 3–5 pulses at 50 Hz repeated at 5 Hz for 40s, a protocol considered to fall into the broader category of ‘repetitive TMS’ (rTMS). Previous studies have shown that the use of rTMS protocols such as cTBS can induce long term changes -- termed by neuroscientists as long-term potentiation (LTP) and long term depression (LTD) -- within M1 or other regions of the brain^8–11^. The changes induced by cTBS are typically monitored and reported by measuring motor evoked potentials (MEPs) and believed to last up to 1 hour post-stimulation^12^. The direction and magnitude of these changes are thought to depend on the pattern of the administered stimulation.^13^ cTBS is thought to induce LTD and therefore suppress neuronal activity after stimulation. While some models have been recognized which describe the electrophysiologic response as a function of stimulation pattern, the precise cellular mechanisms which are at play here are not well understood. Moreover, current efforts to characterize the response to TMS often focus on motor responses, and biomarkers of response to TMS in other areas of the cortex, such as those associated with cognitive processes, have not yet been established.

Current hypotheses include the idea that changes in MEPs are caused by modulation of interneuron and/or pyramidal cell membrane excitability, or reflect a change in synaptic processes^4^. Recent studies have pointed to membrane potentials being influenced primarily at the level of inhibitory/excitatory interneurons within the cortex, as stimulation protocols such as cTBS are generally not considered to be sufficiently intense to activate deeper cortical pyramidal neurons^14^. cTBS specifically is believed to specifically activate inhibitory GABA_A_ and GABA_B_ interneurons within M1 ^12,15,16^. Indeed, multiple studies have found that cTBS and related protocols can induce changes in GABA concentration, as measured by magnetic resonance spectroscopy (MRS) ^17–22^ or have linked TMS-based measures of electrophysiology to GABA content^23,24^. As GABA is known to be the central nervous system’s primary inhibitory neurotransmitter, one account proposes that increased [GABA] may correlate with an inhibitory effect on M1, decreasing measured MEPs.

However, evidence from drug studies suggests that not only the GABA-ergic, but also glutamatergic, systems are perturbed by TMS. For example, the use of glutamatergic NMDA antagonist dextromethorphan has been shown to alter the responses to rTMS protocols^25–27^ and the use of the partial NMDA agonist D-cycloserine has been shown specifically to modulate the effect of theta burst stimulation^28^. A review by Li *et al* has emphasized the role of glutamatergic signaling in a model-based theoretical approach. ^29^ However, despite the apparent consensus that the glutamatergic signaling system is also involved in TMS response, changes in glutamate have not been detected robustly in MRS studies to date of cTBS to M1. It is possible that this is due to the technical limitations (e.g. sensitivity) of the experiments performed to date.

We hypothesized that glutamate, the primary excitatory neurotransmitter, may in fact decrease in M1 upon administration of cTBS. In this study, rather than MRS, we used a more recently developed magnetic-resonance based technique, known as glutamate-weighted chemical exchange saturation transfer or gluCEST^30^, to assess the effects of cTBS on cortical glutamate.

To the best of our knowledge, this technique, or any other which introduces spatial resolution to molecular detection, has not been previously applied to investigate TMS. Given its advantages relative to spectroscopy for measuring glutamate, gluCEST may allow us to better explore the possible role of the excitatory mechanisms in the generation of TMS effects.

GluCEST relies on a different mechanism than spectroscopy: rather than detecting the resonance of the glutamate protons directly, the gluCEST signal originates from the interaction of the glutamate with bulk water, and the signal measured – as in most other forms of MR imaging – is that of the water itself. Specifically, ‘saturation’, refers to a transient decrease in the spin polarization (the origin of the NMR signal) of the protons on the amine group of glutamate by irradiation with a frequency-targeted RF pulse. This saturated population and consequent signal decrease can be transferred to the water by way of the fast chemical exchange occurring between these amine protons and the solvent. In this way, the MR signal and therefore the intensity of the image at any particular point in space reflects the local concentration of glutamate. Note that, because each amine proton position exchanges with water thousands of times per second, the CEST technique provides a corresponding ∼700-fold kinetics-based amplification of the signal-per-mole of protons relative to spectroscopy, in which this ratio is stoichiometric (1:1)^30^. In this work, in addition to benefitting from enhanced sensitivity, we capitalize on the ability of gluCEST to generate spatially resolved brain ‘maps’ weighted for glutamate to investigate the short-term neurometabolic effects of cTBS on healthy humans.

Our study design involves sequential collection of gluCEST images before and after subjects have undergone cTBS or a sham stimulation. Our hypothesis was two-fold: a) the ability to detect small changes in glutamate in a particular location such as M1 would be facilitated by the increased sensitivity of gluCEST; b) the spatial resolution of gluCEST may allow us to observe off-target effects arising either from direct stimulation by residual electric field, or due to internal neural connections themselves.

Our results suggest that we have indeed been able to detect transient shifts in the neurochemical profile of stimulated subjects, both in M1 and adjacent regions, which may underlie the electrophysiological and other more lasting effects attributed to cTBS. This is a pilot study in which only general trends amongst a small population of healthy subjects were determined. However, we believe that this type of data may open doors to further understanding the molecular mechanisms of TMS, the geometric and temporal extent of its effects, the varying effect of different TMS protocols, and the origin of variability in response to TMS within the population.

## Results

GluCEST imaging was performed on fifteen healthy subjects before and after receiving either a real (n= 10) or sham (n = 5) cTBS session (**Figure 1**). To facilitate accurate slice placement, the left M1 was identified in-magnet by use of BOLD with real-time processing; the data was then subjected to regional analysis *post facto* using segmentation by Freesurfer. Visual comparison of the segmentation results (Figure 2A, red) with the BOLD analysis (Figure 2B) confirms the success of the in-magnet localization of M1 for axial slice placement (Figure 2C).

**Figure 1.**
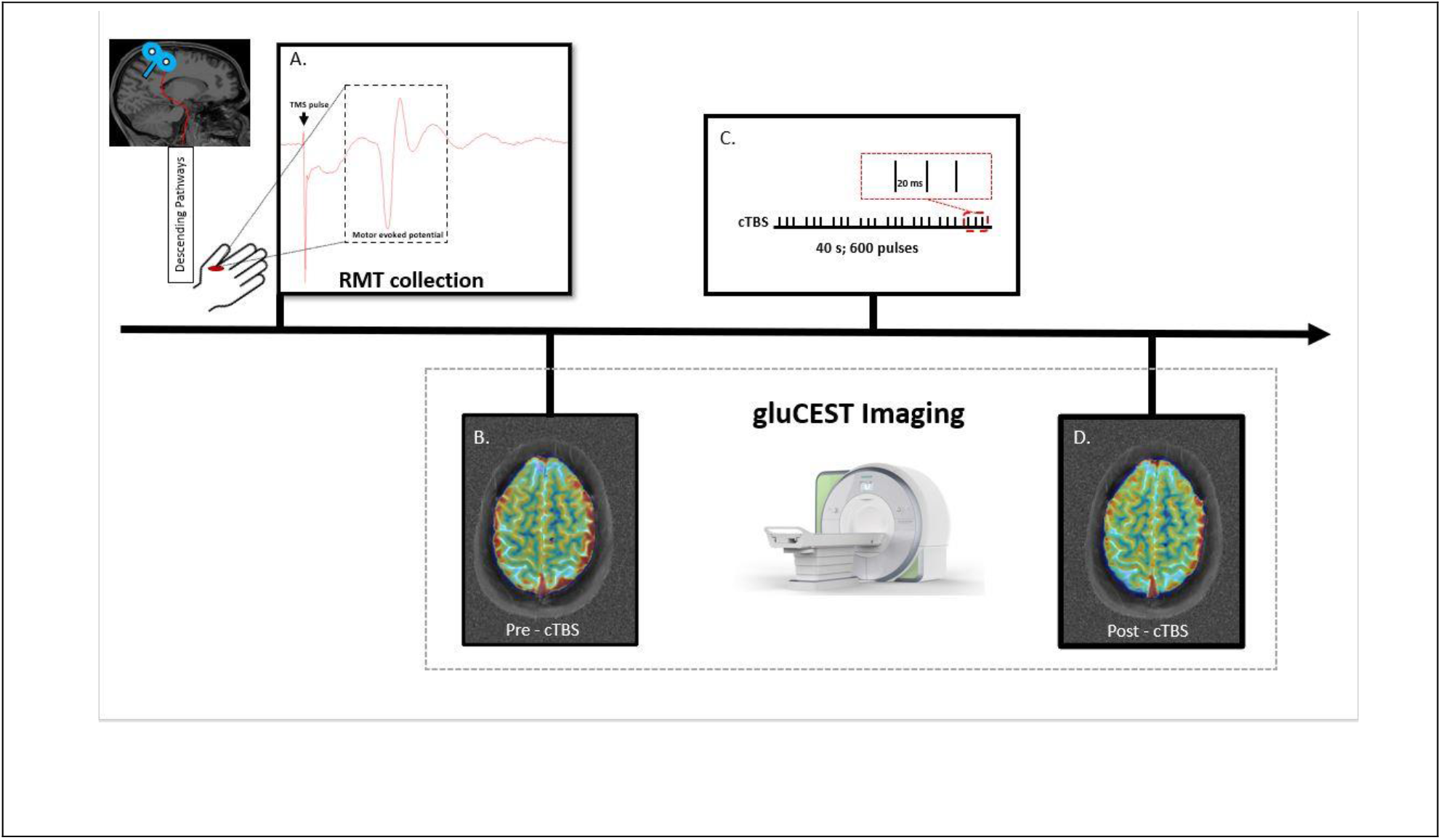
Overview of TMS/gluCEST experiment. **A**. Prior to initial imaging, subjects undergo TMS to determine resting motor threshold (RMT). **B**. An initial gluCEST map is acquired in a slice that includes the motor cortex. **C**. Subjects are removed from the magnet and receive cTBS to the left motor cortex in accordance with the illustrated pattern. Some subjects receive sham stimulation. **D**. A second gluCEST map is acquired after stimulation or sham.

**Figure 2.**
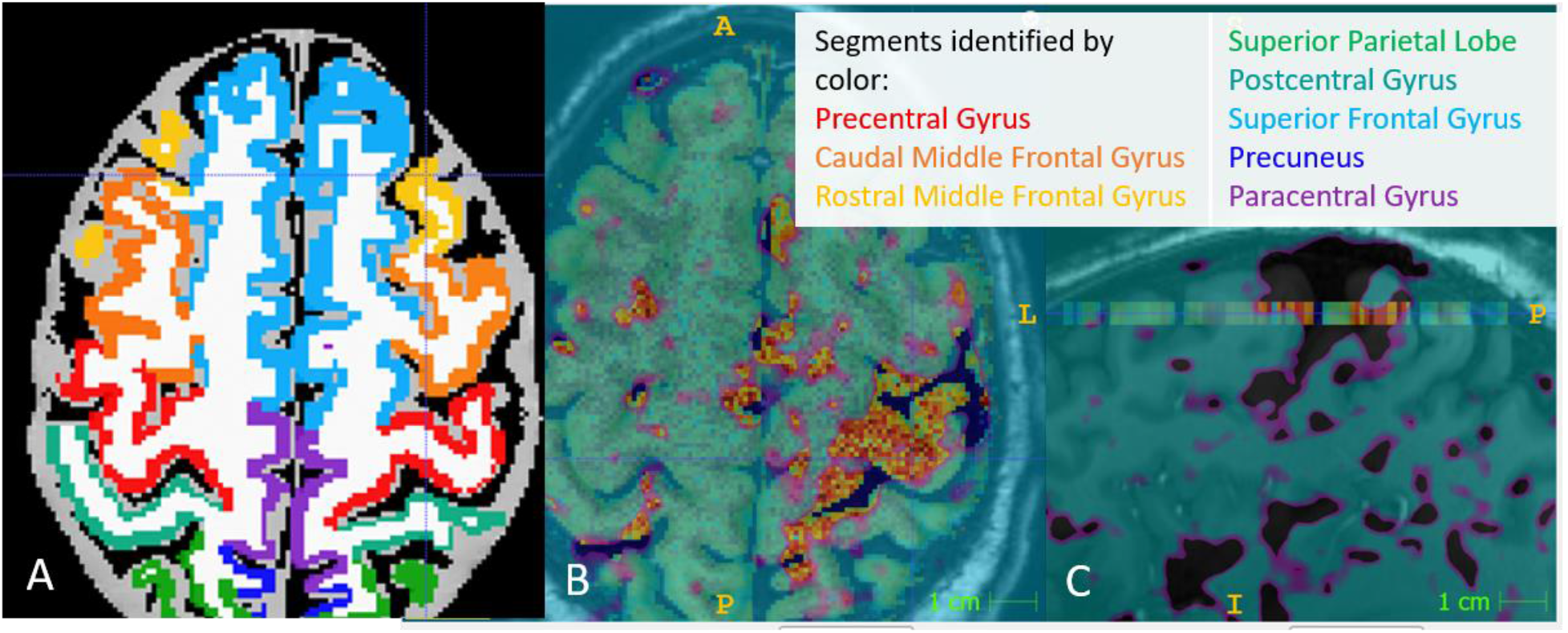
Illustration of CEST slice placement and regional analysis. A) CEST slice shown as segmented by Freesurfer’s ReconAll. The precentral gyrus is shown in red. B) View of CEST slice overlaid with structural image and BOLD results used to localize this slice before imaging experiment. The transparent area represents the region identified by BOLD as the targeted location. Good correspondence is observed between this real-time functional measurement and the atlas-based segmentation performed *post facto*. C) Same image overlay series as B, shown in sagittal view: CEST slice after post-processing (color) shown intersecting the black/transparent BOLD-identified ROI.

An example of gluCEST data masked for the precentral gyrus (local of M1) from one stimulated subject with a visually apparent change in this region is shown in **Figure 3**, along with the corresponding histograms of the precentral gyrus pixel-wise gluCEST data pre and post-cTBS. A slight left-shift can be observed in the center of the histogram, representing a decrease in the median gluCEST value. When analyzing data from all subjects, the 99% confidence interval for the change in gluCEST in the precentral gyrus upon stimulation was [-.11--.33] % asymmetry. The average pre-stimulation gluCEST value for left precentral gyrus was 8.3% asymmetry. Thus, the values computed in this confidence interval represent a [1.3- 4%] decrease in the gluCEST signal. No statistically significant change was detected in the precentral gyrus in the sham subjects (see **Table 1**). **Figure 4** shows the average gluCEST changes over the precentral gyrus ROI for the right and left sides for all subjects. No trend is apparent in the right precentral gyrus, or in the left (ipsilateral) hemisphere of the sham subjects. However, eight out of ten stimulated subjects exhibited an ROI average decrease in gluCEST in the precentral gyrus upon receiving cTBS.

**Table 1.**
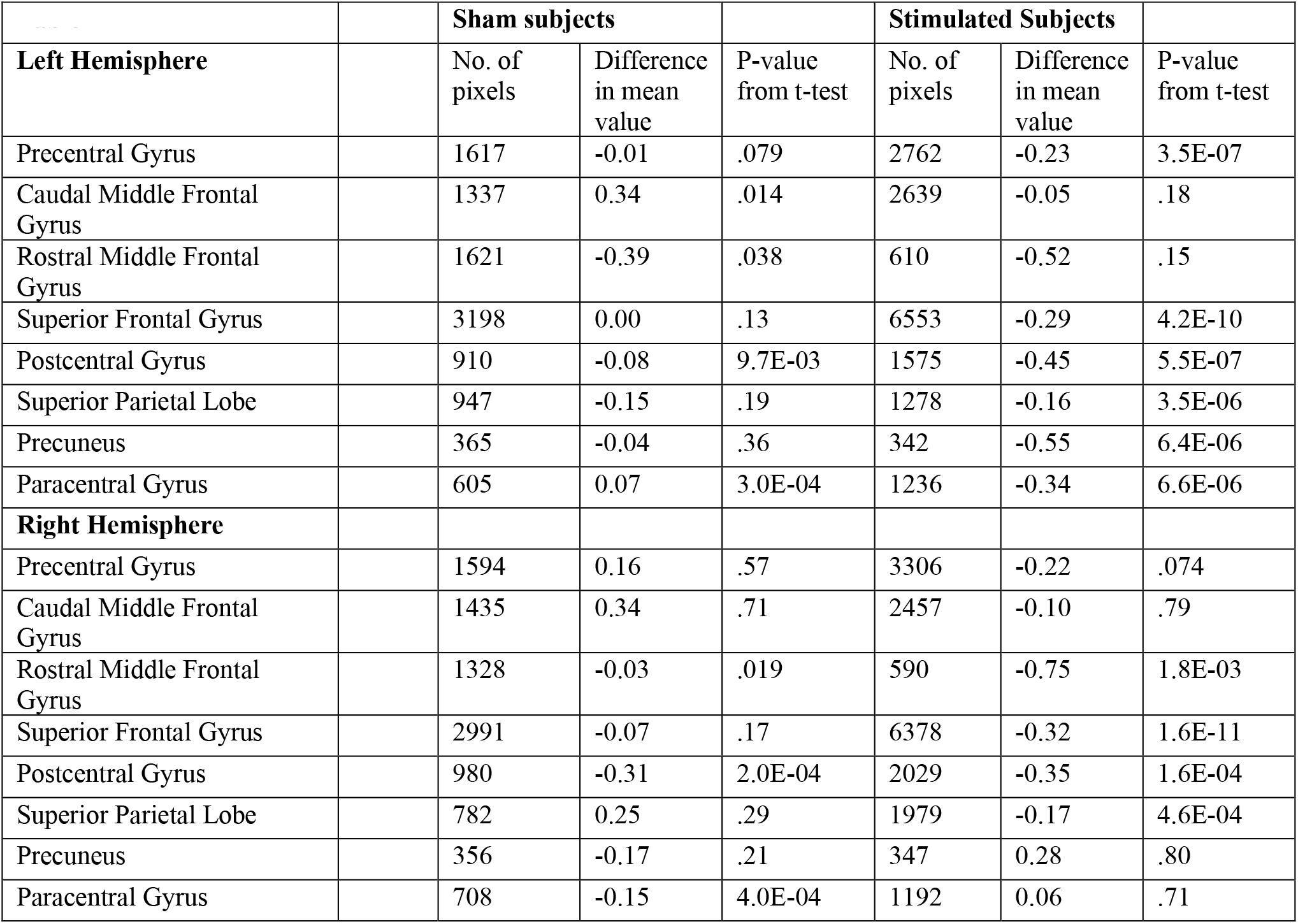

| **Table 1** |  | **Sham subjects** |  |  | **Stimulated Subjects** |  |  |
| --- | --- | --- | --- | --- | --- | --- | --- |
| **Left Hemisphere** |  | No. of pixels | Difference in mean value | P-value from t-test | No. of pixels | Difference in mean value | P-value from t-test |
| Precentral Gyrus |  | 1617 | -0.01 | .079 | 2762 | -0.23 | 3.5E-07 |
| Caudal Middle Frontal Gyrus |  | 1337 | 0.34 | .014 | 2639 | -0.05 | .18 |
| Rostral Middle Frontal Gyrus |  | 1621 | -0.39 | .038 | 610 | -0.52 | .15 |
| Superior Frontal Gyrus |  | 3198 | 0.00 | .13 | 6553 | -0.29 | 4.2E-10 |
| Postcentral Gyrus |  | 910 | -0.08 | 9.7E-03 | 1575 | -0.45 | 5.5E-07 |
| Superior Parietal Lobe |  | 947 | -0.15 | .19 | 1278 | -0.16 | 3.5E-06 |
| Precuneus |  | 365 | -0.04 | .36 | 342 | -0.55 | 6.4E-06 |
| Paracentral Gyrus |  | 605 | 0.07 | 3.0E-04 | 1236 | -0.34 | 6.6E-06 |
| **Right Hemisphere** |  |  |  |  |  |  |  |
| Precentral Gyrus |  | 1594 | 0.16 | .57 | 3306 | -0.22 | .074 |
| Caudal Middle Frontal Gyrus |  | 1435 | 0.34 | .71 | 2457 | -0.10 | .79 |
| Rostral Middle Frontal Gyrus |  | 1328 | -0.03 | .019 | 590 | -0.75 | 1.8E-03 |
| Superior Frontal Gyrus |  | 2991 | -0.07 | .17 | 6378 | -0.32 | 1.6E-11 |
| Postcentral Gyrus |  | 980 | -0.31 | 2.0E-04 | 2029 | -0.35 | 1.6E-04 |
| Superior Parietal Lobe |  | 782 | 0.25 | .29 | 1979 | -0.17 | 4.6E-04 |
| Precuneus |  | 356 | -0.17 | .21 | 347 | 0.28 | .80 |
| Paracentral Gyrus |  | 708 | -0.15 | 4.0E-04 | 1192 | 0.06 | .71 |

**Figure 3:**
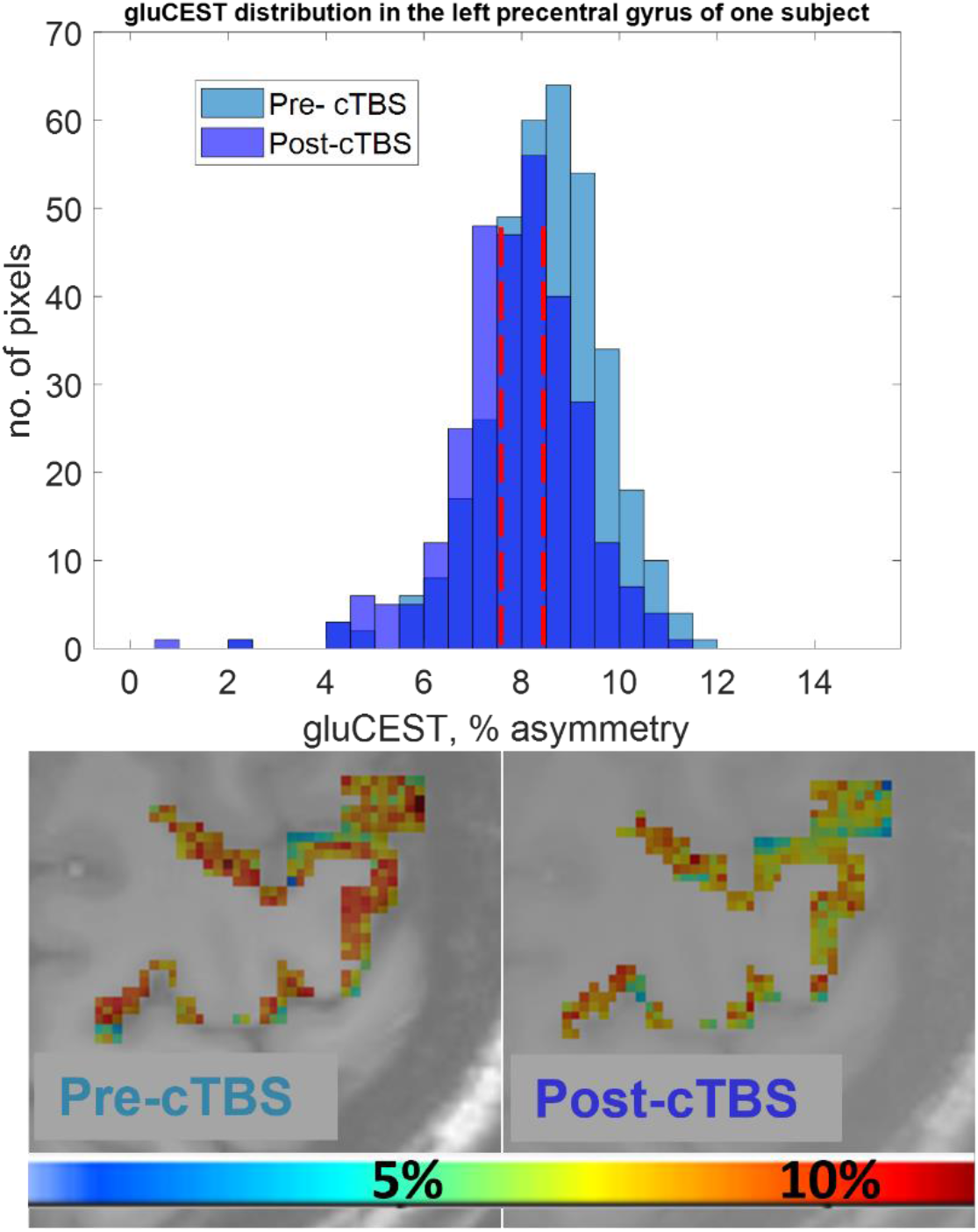
Masked CEST maps and histogram for gluCEST in the left precentral gyrus of one stimulated subject. Color scale in CEST maps is 0-13%, negative normalized asymmetry. The slight left shift in the histogram reflects the color change seen in the maps. The 99% confidence interval for the mean change in the left precentral gyrus (locus of M1) for all stimulated subjects was .13-.33% gluCEST contrast. In the subject shown, the change in mean gluCEST was .88% gluCEST contrast. Red bars indicate the mean gluCEST before and after cTBS in this subject (the mean has a slightly lower value than the median in each distribution).

**Figure 4:**
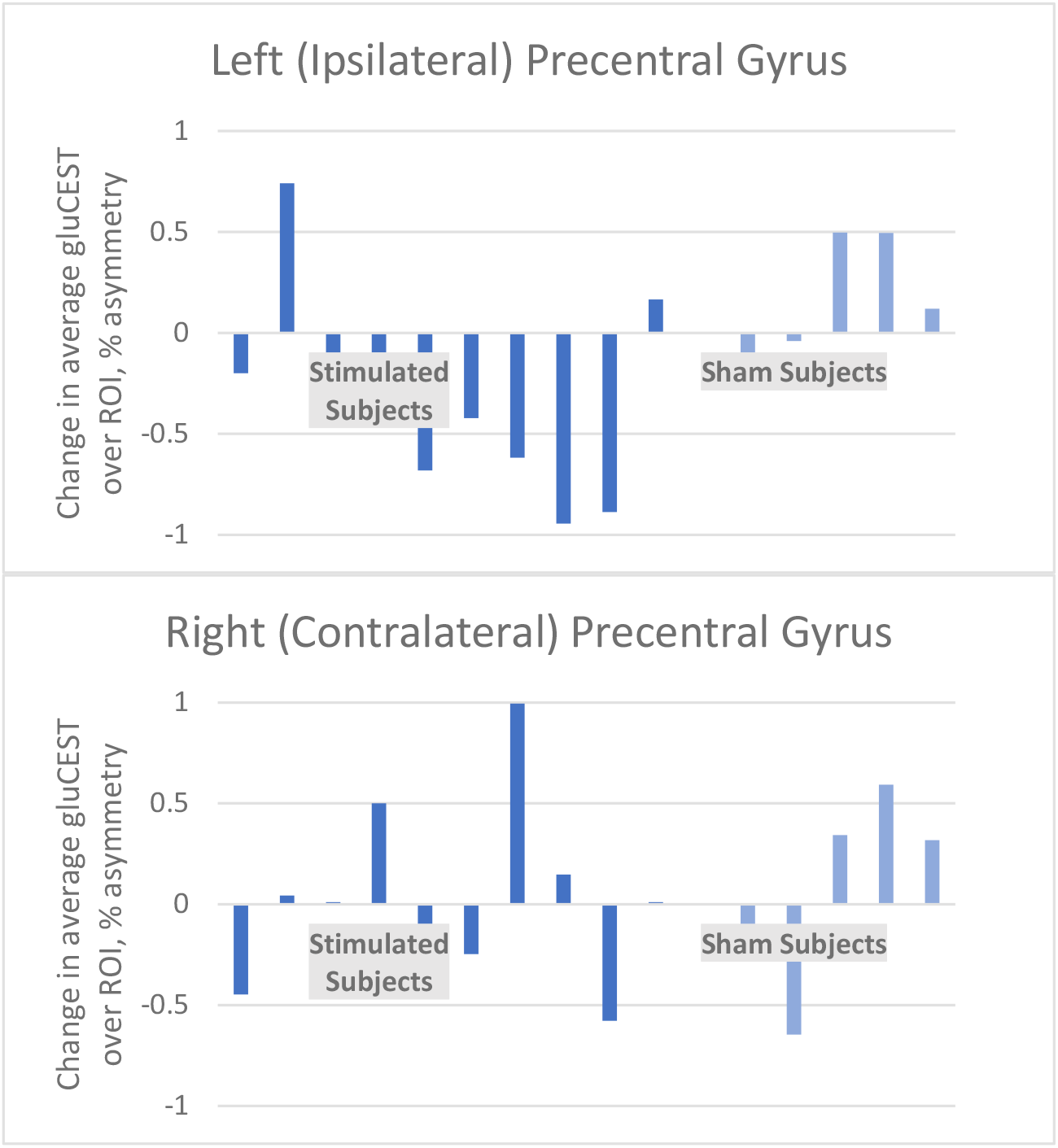
Barplots of average change in gluCEST signal, in right and left precentral gyri (locus of M1), displayed by individual subject. For stimulated subjects, n = 10; sham, n = 5.

While initial analysis focused on M1, a decrease in gluCEST signal throughout the left (ipsilateral) hemisphere was detected in subjects who received cTBS. **Figure 5** shows segment-wise average gluCEST data from all subjects, projected onto the anatomy of a single subject for visualization. **Figure 5A** shows the average gluCEST values by segment in the baseline scan, containing data from all 15 subjects before they had received either the sham or real stimulation. Average gluCEST values range from 7-9%, typical for gray matter regions in healthy subjects using this protocol^31,32^. Interestingly, there appears to be some anatomic variability in gluCEST values in this slice at baseline; however, we did not attempt to further interpret these results, given the limited number of subjects. **Figure 5B** shows the same data in the sham subjects (n = 5), ‘post-sham’. These values remain largely unchanged from the baseline values for the whole group shown in **5A**. As shown in **Figure S3**, the confidence intervals for all segments in sham subjects span zero and lack statistical significance (also see Table 1). In contrast, **Figure 5C**, showing the post-stimulation (cTBS) gluCEST averages (n = 10), is significantly different from the baseline and post-sham results. Several segments, not only M1 (stimulation target, indicated by green arrow) show a significant decrease in average gluCEST value compared to the baseline measurement.

**Figure 5.**
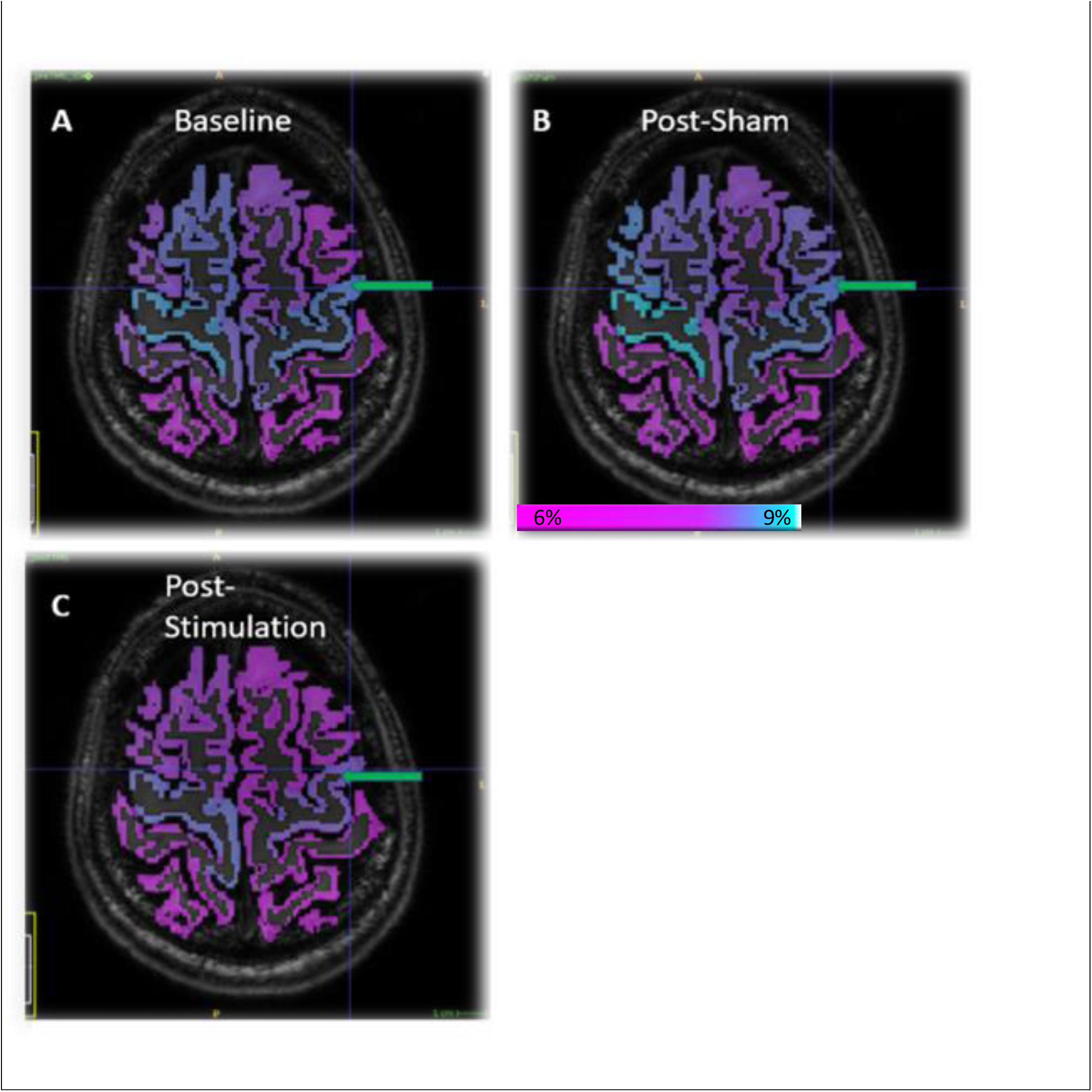
Segmentwise gluCEST maps. Data from all subjects projected onto the anatomy of a single subject for visual representation. A) average gluCEST by segment, baseline (pre-stimulation) −15 subjects. B) post “sham” (placebo) stimulation - 5 subjects. C) post cTBS (real) stimulation -- 10 subjects. The colorscale is identical in all maps, 6-9% gluCEST contrast, as indicated by the colorbar. Green arrow indicates the left precentral gyrus, the intended target of cTBS in stimulated subjects. Please see the color coding in Figure 4.1 for full listing of anatomical segments treated distinctly in this analysis.

For statistical analysis, unpaired T-tests were performed comparing the pixelwise data of these segments between the ‘pre’ and ‘post’ scans for both groups. **Table 1** lists the total number of pixels in the distribution representing each ROI, the mean over those pixels, and the corresponding P-value of the T-test which compared the pre-cTBS and post-cTBS distributions. Analyzed in this way, statistically significant changes (decreases) in gluCEST were found in many segments in the left hemisphere of the brain of stimulated subjects. No P-values less than 1e-4 were calculated for sham subject data. Interestingly, the left precentral gyrus – the locus of M1—is not the segment with the greatest change in mean value, and the contralateral right precentral gyrus is not amongst those segments in which a strongly significant change was observed. The confidence intervals from these T-tests for the change upon stimulation or sham are visualized in barplots in **Figure S3**. Note that T-tests were unpaired, as imperfect registration between scans prohibits a 1:1 correlation of every pixel in the ROI.

## Discussion

In this study, we used gluCEST, a form of MRI in which the image contrast is weighted for the presence of glutamate, to image the brains of healthy volunteers who have undergone cTBS or a sham protocol. We find that on average, statistically significant changes appear in the gluCEST values from the brains of stimulated subjects; specifically, that gluCEST decreases in M1 and other regions of the ipsilateral hemisphere in stimulated subjects, but not in the contralateral hemisphere, and not in subjects who received a ‘sham’ stimulation between imaging sessions. The gluCEST decrease was on the order of 0-5% of the baseline value (varying between subject and area of the brain). There are perhaps two separate and interesting observations here which merit discussion and interpretation in the context of existing literature: a) that gluCEST decreased, as opposed to increasing or remaining unchanged, upon administration of cTBS and b) the spatial distribution of the measured change.

According to the review by Cuypers and Marsman^33^, a number of apparently inconsistent results have been reported regarding the existence and sign of a correlation between TMS-based measures of electrophysiology and MRS-based measurements of glutamate and other metabolites. For example, one study found a positive correlation between the concentration of glutamate in M1 and the MEP input/output slope^34^, while another study found a negative correlation between this measurement and the plateau of the MEP input/output curve^35^. With regard to glutamate, such inconclusive results are not surprising, given the limitations of spectroscopy for detecting such changes, both in terms of sensitivity and spectral resolution from glutamine. Moreover, variability in the TMS protocols under investigation adds degrees of freedom that make such meta-analysis difficult.

The TMS study of Dyke *et al* 2017 was of the few to perform spectroscopy at 7T, where differentiation between glutamate and glutamine becomes possible^35^. They found statistically significant correlations between spectroscopic measurements of glutamate and two different TMS-based measures: quantities described as “Intracortical Facilitation” and the “Input/Output plateau”. Interestingly, they find that these two correlations are of opposite sign, suggesting that the relationships between metabolite concentrations and various electrophysiologic parameters may be more complex and sensitive than many of the existing analyses suggest. Our measurement indicates that the concentration of glutamate decreases in the brains of stimulated subjects at ∼30 minutes post-cTBS. It is difficult to say definitively whether, on the whole, this imaging result corroborates existing work about cTBS physiology, as no directly comparable experiment has been done before.

The observation of the spatial distribution of a TMS-induced metabolic change other than BOLD is unique to this study. GluCEST is the only way to detect glutamate *in vivo* with spatial resolution comparable to other forms of magnetic resonance imaging (MRI). While our results measuring the spatial distribution of the TMS-induced changes are preliminary, they serve to highlight that the TMS pulse effects more than just its nominal target, which is often a target in the primary motor cortex whose stimulation results in an observable muscle motion and MEP. The spatial distribution of the effects suggests that off-target responses may have more to do with neural connectivity than residual strength of the pulsed field itself. Interestingly, the only contralateral effect observed was in the superior frontal gyrus. We anticipate additional informative findings about the spatial profile of TMS-induced metabolic effects as we replace the single-slice gluCEST image with a larger, volumetric acquisition in future studies.

In conclusion, we believe that the increased sensitivity of gluCEST relative to spectroscopy has allowed us to detect subtle changes in glutamate concentration that so far have been inaccessible to TMS researchers relying on single-voxel MRS. Furthermore, our results provide information about the spatial distribution of the TMS effect which is unprecedented in existing measurements. Further studies should examine if gluCEST imaging can be used to associate glutamate changes with changes in cognition, which would provide a needed biomarker for cognitive changes within TMS studies. We refrain at this time from putting forth any specific mechanistic conclusions about cTBS based on our observations, as the goal of this preliminary study was simply to explore the utility of gluCEST as a method to study TMS. The presence of coherent, statistically significant findings despite the small number of participants in the study is very encouraging, and illustrates that indeed the spatially resolved molecular imaging capabilities of gluCEST have a role to play in elucidating the chemical mechanisms of therapy by non-invasive brain stimulation. Moreover, it could be possible to obtain reliable subject-specific biomarkers that predict neurotransmitter-mediated TMS outcomes. The obvious areas for expansion in our further work include implementation of a newly developed volumetric (3D) gluCEST protocol and, in an effort to produce results which can be more directly compared with other studies, acquisition of additional types of MEP-related measures.

## Materials and Methods

### Human subjects information

15 healthy individuals (*mean age* = 29.5, *SD*. = 8.8 years, 5 female) were recruited for this study. Subjects provided informed, written consent and all procedures were carried out in accordance with the guidelines of the Institutional Review Board/Human Subjects Committee at University of Pennsylvania. TMS was administered to all subjects in accordance with the procedure described in below. Subjects were randomly assigned to either the active stimulation group (n = 10) or the sham group (n = 5). All subjects were right-handed participants and had no prior history of neurologic or psychiatric disease.

### cTBS and sham

The TMS coil was used to find the subject’s contralateral right hand first dorsal interosseous (FDI) muscle via the neuronavigation software Brainsight (Rogue Research, Montreal, Québec, Canada) and the subject’s T1w image. A subject’s resting motor threshold (RMT) was determined when the FDI muscle was activated at rest 50% of the time, as determined by motor evoked potentials (MEPs) at a peak-to-peak amplitude of 1 mV. A subject’s active motor threshold was determined when the FDI muscle was engaged in a motor task 50% of the time (determined by MEPs) while the subject maintained a 20% maximum voluntary contraction of the muscle using a visual feedback monitor. The resting and active motor threshold values (i.e. the machine output when at rest or doing the active task) were noted. The TMS coil was held at an approximately 45-degree angle relative to the subject’s precentral gyrus. Subjects then participated in two imaging sessions (“pre-TMS” and “post-TMS”). Between the two sessions, subjects were randomized to either active or sham TMS. In the active TMS group, we administered continuous theta burst stimulation^12^ (cTBS), to the FDI muscle target on the motor cortex. cTBS is a commonly used neural stimulation protocol which consists of busts containing 3 pulses at 50Hz and an intensity at 80% of the subject’s active motor threshold (AMT). The bursts are repeated continuously for a total of 40s and 600 pulses (**Figure 1c**). Stimulation was delivered with a 70mM hand-held figure-eight coil at 45-degrees from the subject’s precentral gyrus for the duration of the 40s of cTBS using a Magstim Super Rapid Stimulator (Magstim, Inc., Whitland, Dyfed, UK.).

### MRI acquisition procedure

All images were obtained on a Siemens 7.0T MAGNETOM Terra scanner (Siemens Healthcare, Erlangen, Germany) outfitted with a single volume transmit/32 channel receive phased array head coil (Nova Medical, Wilmington, MA, USA). All subjects underwent two sessions of MR imaging: one prior to either cTBS or sham stimulation, and one directly following the stimulation session. Between-session slice registration was accomplished using the in-house program *ImScribe*, described previously^36^. Prior to beginning the initial CEST measurement, structural and BOLD scans (fMRI) were acquired and processed in order to locate the target region of stimulation for accurate placement of the CEST slice. Subjects were instructed to perform voluntary motion of the first dorsal interosseus muscle (index finger), the same motion which is induced involuntarily by the cTBS protocol when administered above motor threshold. This voluntary motion led to a localized BOLD signal that could be visualized and then registered to the structural image visible on the scanner interface where slice placement is performed by the operator. The CEST slice was then acquired in such a fashion as to maximally capture the activated region. See Figure 4.1 for illustration.

GluCEST data was collected with a single-slice CEST sequence based on gradient-recalled echo with the following parameters: TR/TE = 4.7/2.3 ms, 10° flip angle, 5 mm slice thickness, with 0.75 × 0.75 mm2 in plane resolution over a 156 × 192 mm field of view. Magnetization preparation was achieved using eight 3.1µT RMS amplitude, 98 ms Hamming-window shaped pulses with 2 ms inter-pulse delay applied at offset frequencies {±1.8, 2.1, 2.4, 2.7, 3.0, 3.3, 3.6, 3.9, 4.2} ppm relative to water. Additional acquisitions over the same field of view included a water saturation acquisition (WASSR) for B_0_ mapping^37^, a flip/crush sequence for B_1_ mapping^38^, and the Siemens product sequence MP2RAGE for generation of T_1_ maps. A ‘reference’ image consisting of only the read-out module of the CEST sequence (with no saturation) was also collected for each slice. Please note that ‘% asymmetry’, referring to the difference between the signal arising from saturated and non-saturated signals, is a standard unit for gluCEST measurement.

### Data processing and analysis

CEST-weighted images were corrected for the B_0_ field distribution using the B_0_ image generated by the WASSR scan, as described in ^30^. CEST images were corrected for B_1_ inhomogeneity using a recently developed procedure based on B_1_ and T_1_ mapping^32^. T_1_ and T_2_ weighted full-brain structural images were used for segmentation by Freesurfer’s ‘Recon All’ function^39^ (Freesurfer: Martinos Center for Biomedical Imaging, Charlestown, MA, USA). The output from Recon All (segmentation image) was transformed back into the original acquisition space, and resliced to correspond to the CEST acquisition for pixelwise regional analysis. Regional averages and distributions were calculated and visualized using in-house code written in Matlab (Mathworks, Natick, MA, USA); statistical analysis was performed using Matlab’s unpaired T-test function. Plots and visualizations were generated using Matlab and ITK-SNAP^40,41^.

## Acknowledgements

The authors would like to thank the numerous subjects who volunteered to participate in this study, particularly during lengthy pilot experiments.

## Author contributions

**A.T.J.C**. Methodology, Software, Formal Analysis, Investigation, Writing-Original Draft

**B.L.D**. Methodology, Formal Analysis, Investigation, Writing-Editing and Review

**J.P.Z**. Methodology

**A.K**. Investigation, Resources

**B.E**. Investigation, Methodology

**O.F**. Investigation, Resources

**M.A.E**. Methodology, Software

**R.H.H**., **H.B.C**. Supervision, Manuscript Preparation

**R.R**. Conceptualization, Supervision

**J.D.M**. Conceptualization, Supervision, Manuscript Preparation

