## Supplemental Figures for "Glutamate-Weighted Magnetic Resonance Imaging (GluCEST) Detects Effects of Transcranial Magnetic Stimulation to the Motor Cortex"

**Supplementary Materials:**

S1) Flowchart of gluCEST acquisition and post-processing (reproduced from ref. 32)
S2) Example full-slice gluCEST maps
S3) Barplot of confidence intervals of gluCEST changes
S4) Subject-wise plots of mean gluCEST change by segment

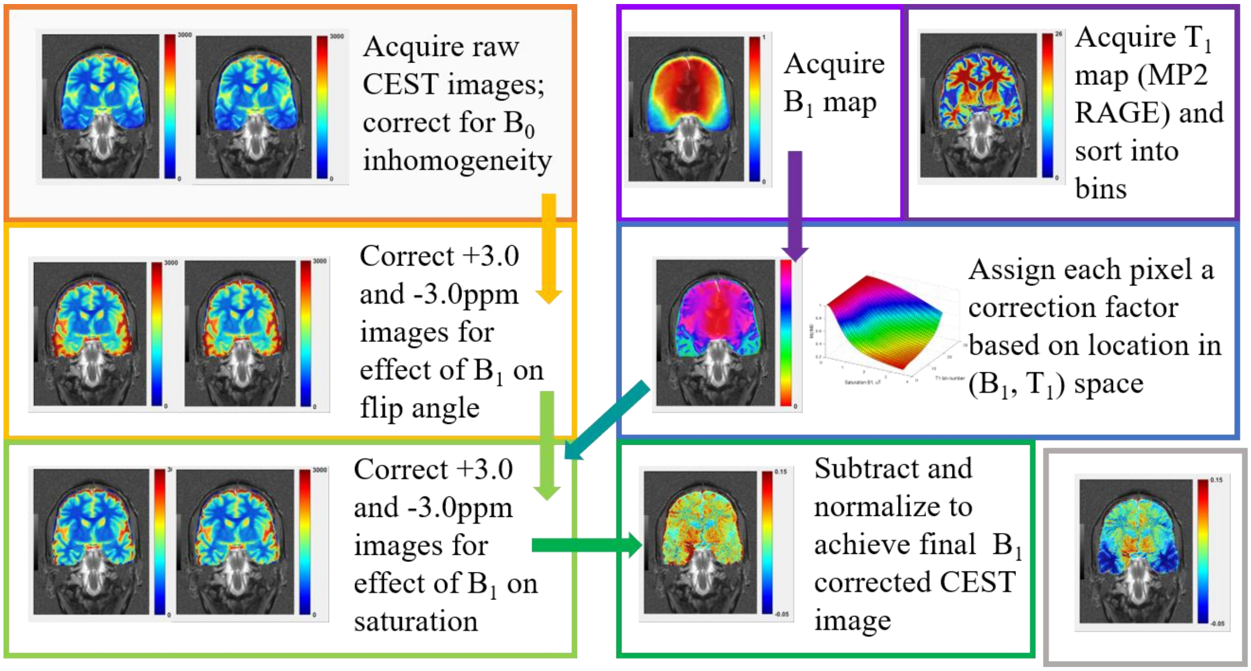


**Figure S1. Schematic of gluCEST acquisition and post-processing using $B_1$ correction based on $M_Z(B_1, T_1)$ surfaces [32].** The top row of boxes represents images acquired directly or with additional, existing post - processing steps during the experiment: raw CEST images (then subject to $B_0$ correction based on WASSR acquisition); $B_1$ map (based on flip/crush acquisition); $T_1$ map generated by MP2RAGE. Subsequent steps proceed as indicated by arrows, to arrive at the final output, a fully "$B_1$ corrected" gluCEST image (green box). The same example image with no correction for $B_1$ inhomogeneity is shown in the gray box. Please note that the images provided in this schematic are not intended for close inspection or evaluation, but are intended only to add concreteness and aid the reader in following the description of the process.

**Figure S2**. **Example full-slice gluCEST images.** Pre and post images for two stimulated subjects (top, middle rows) and one sham subject (bottom row) are shown. Color map range is 0-15% negative-normalized asymmetry. GluCEST images for the brain ROI are shown unfiltered (in original resolution) and overlaid with the $T_1$-weighted structural image.

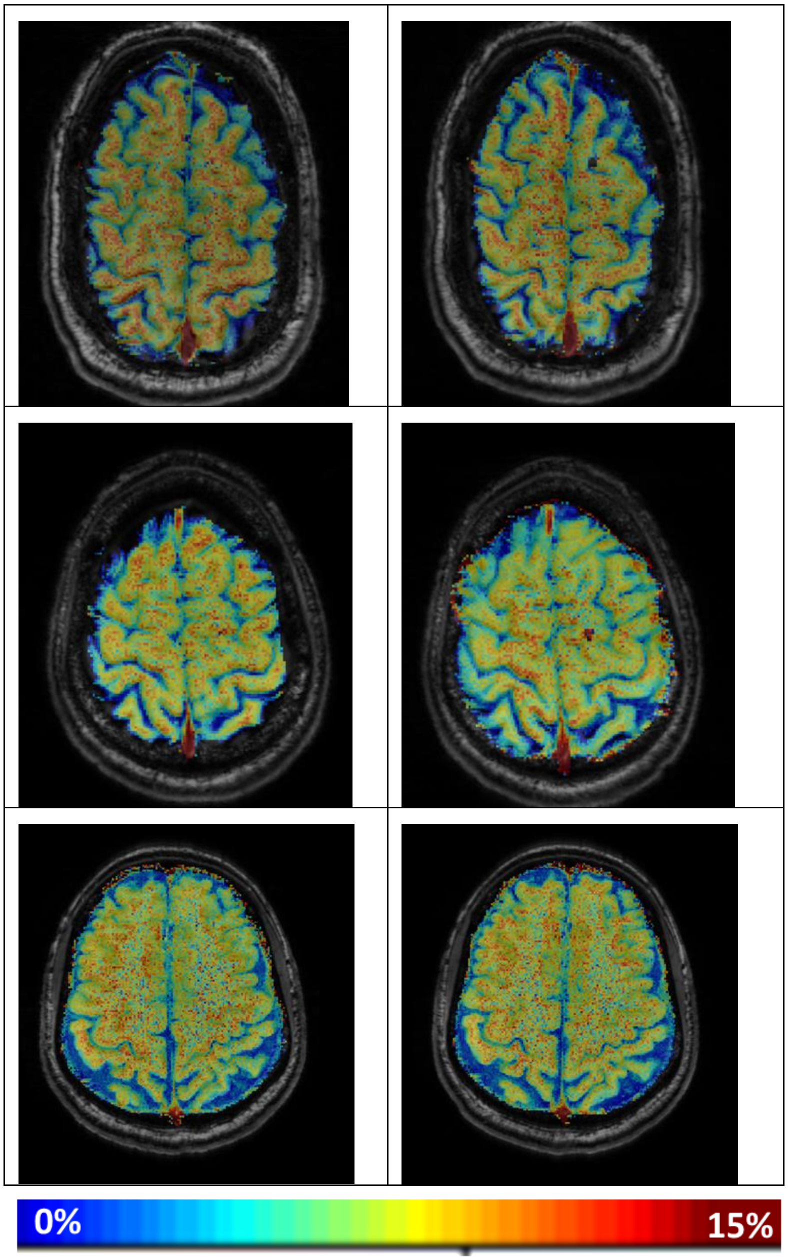


| | |
|---|---|
| Pre- cTBS (top two rows)<br>Pre-sham (bottom row) | Post- cTBS (top two rows)<br>Post-sham (bottom row) |

**Figure S3:** Barplot visualization of 99% confidence intervals (CI) of mean change, as reported by unpaired T-test (enumerated in Table 1). A cluster of two bars, representing the upper and lower bound of the confidence interval, is shown for each segment, beginning with the Precentral Gyrus for each side. The 'true' change likely lies between the two confidence intervals, generally giving a value near zero for sham subjects and most right hemisphere segments, but a small negative value for stimulated subjects. The double asterisks indicate where p-values reflected a high statistical significance of the change.

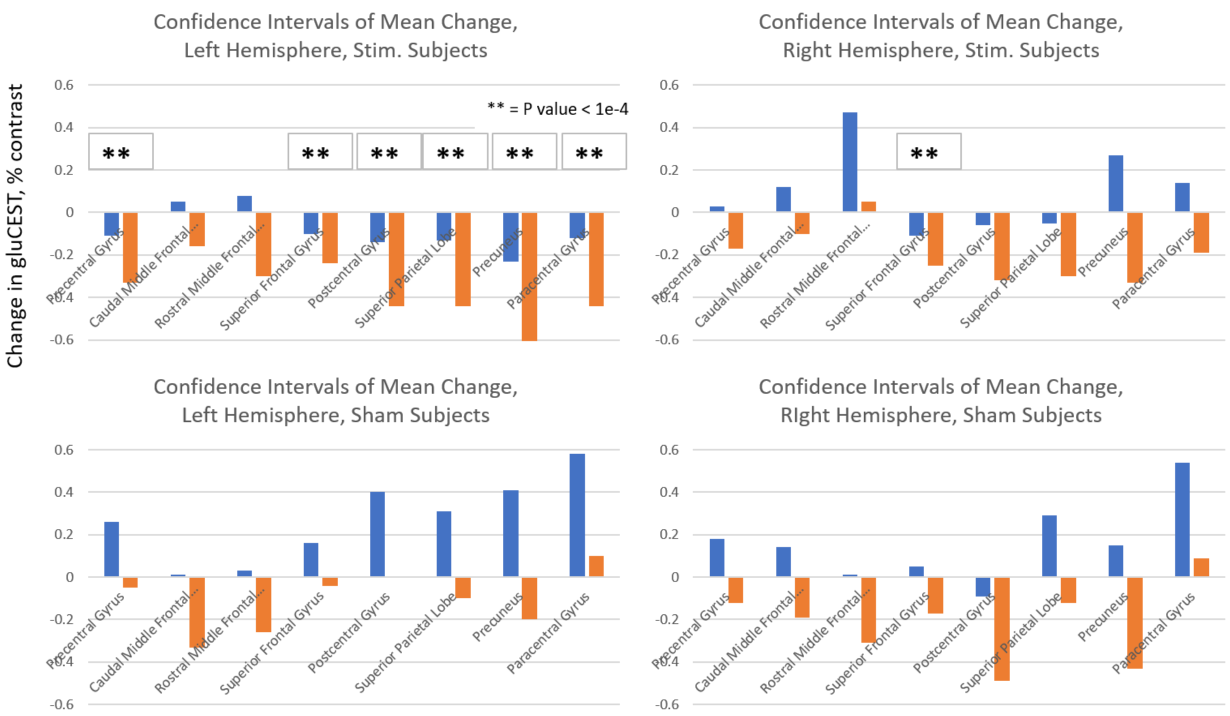

**Figure S4:** Subject-wise plots of mean gluCEST change over ROI for remaining anatomical segments included in the analysis (analogous to the data shown for left and right Precentral Gyrus in Figure 4).

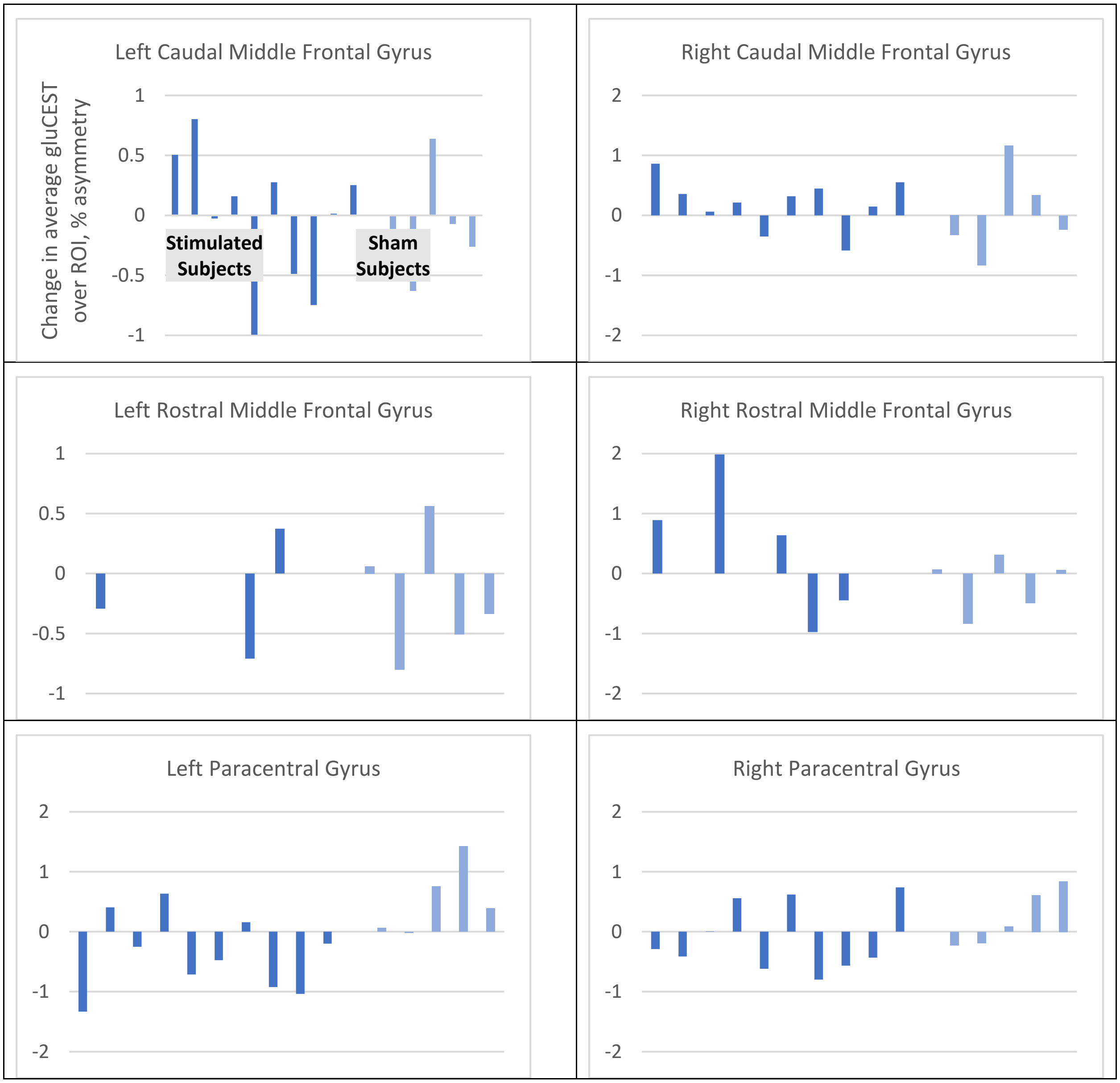

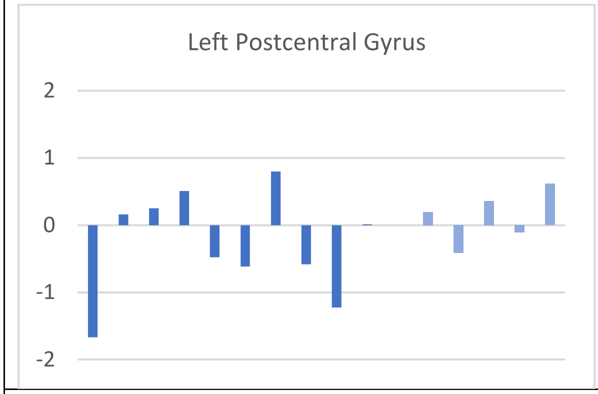
Left Postcentral Gyrus
2
1
0
-1
-2

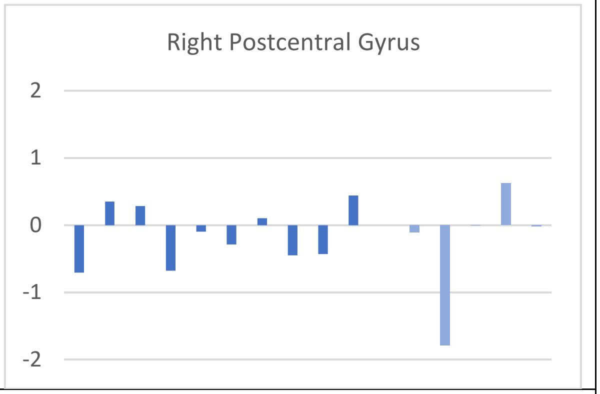
Right Postcentral Gyrus
2
1
0
-1
-2

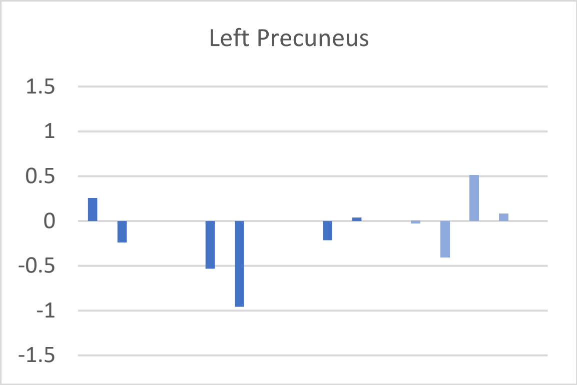
Left Precuneus
1.5
1
0.5
0
-0.5
-1
-1.5

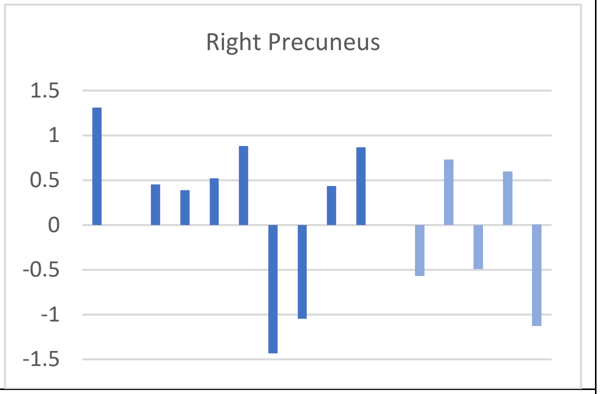
Right Precuneus
1.5
1
0.5
0
-0.5
-1
-1.5

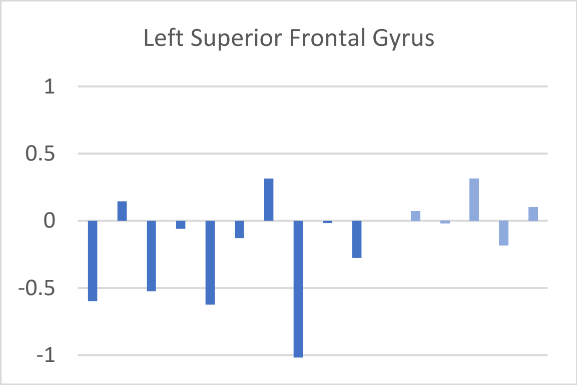
Left Superior Frontal Gyrus
1
0.5
0
-0.5
-1

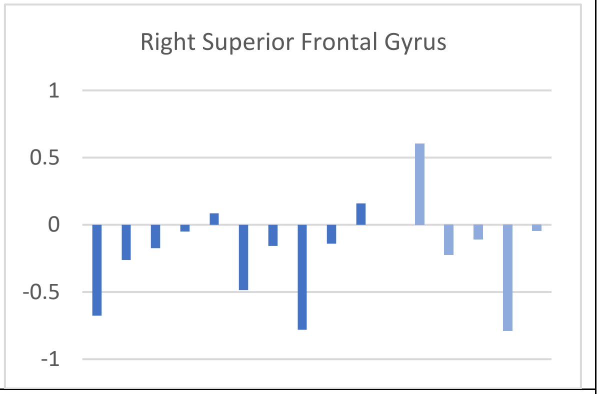
Right Superior Frontal Gyrus
1
0.5
0
-0.5
-1

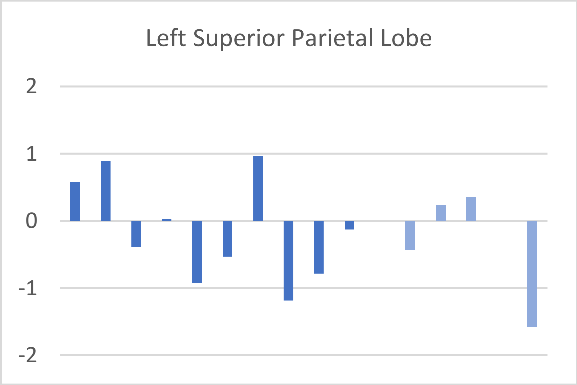
Left Superior Parietal Lobe
2
1
0
-1
-2

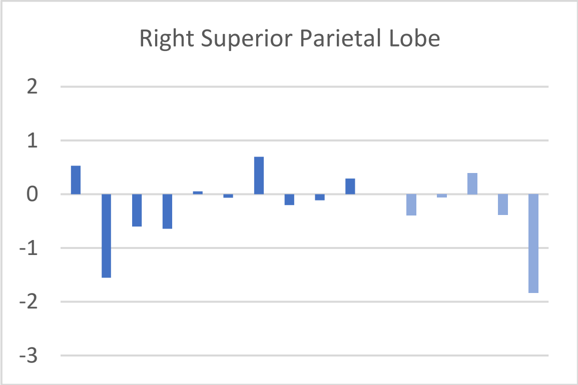
Right Superior Parietal Lobe
2
1
0
-1
-2
-3
